# Speech Production in Intracranial Electroencephalography: iBIDS Dataset

**DOI:** 10.1101/2022.04.05.487183

**Authors:** Maxime Verwoert, Maarten C. Ottenhoff, Sophocles Goulis, Albert J. Colon, Louis Wagner, Simon Tousseyn, Johannes P. van Dijk, Pieter L. Kubben, Christian Herff

## Abstract

Speech production is an intricate process involving a large number of muscles and cognitive processes. The neural processes underlying speech production are not completely understood. As speech is a uniquely human ability, it can not be investigated in animal models. High-fidelity human data can only be obtained in clinical settings and is therefore not easily available to all researchers. Here, we provide a dataset of 10 participants reading out individual words while we measured intracranial EEG from a total of 1103 electrodes. The data, with its high temporal resolution and coverage of a large variety of cortical and sub-cortical brain regions, can help in understanding the speech production process better. Simultaneously, the data can be used to test speech decoding and synthesis approaches from neural data to develop speech Brain-Computer Interfaces and speech neuroprostheses.

## Background & Summary

Brain-Computer Interfaces (BCIs^1^) that directly decode speech from neural activity have recently gained large attention as they could provide an intuitive means of communication for patients who lost the ability to speak^2–7^.

The creation of a speech neuroprosthesis depends on a firm understanding of the speech production process in the brain, the particular timing of brain regions involved and where to best decode them. Despite a number of existing models on the speech production process^8, 9^, the precise role of all areas involved has yet to be understood. Recent advances highlight that deeper brain structures, such as the hippocampus^10–12^ and thalamus^13, 14^, are also involved in language in general and speech production specifically. A dataset providing accessible data for a simple speech production task in cortical and deeper brain structures could help to further understand this intricate process.

Despite the fact that a full understanding of speech production is currently lacking, great advances have been made in the field of speech neuroprostheses recently. The decoding of a textual representation by decoding phonemes^15, 16^, phonetic^17^ or articulatory^18^ features, words^19^, full sentences^20–23^ or spotting of speech keywords^24^ is possible from neural recordings during actual speech production. Results are becoming robust enough for first trials in speech impaired patients^7^. To facilitate more natural communication, some studies aimed at directly synthesizing an audio waveform of speech from neural data recorded during speech production^25–28^. Initial results indicate that the decoding of speech processes is possible from imagined speech production from offline data^29–31^ and in real-time^32, 33^.

Most of these recent advances employ electrocorticography (ECoG), an invasive recording modality of neural activity that provides high temporal and spatial resolution and high signal-to-noise ratio^34^. Additionally, ECoG is less affected by movement artifacts than non-invasive measures of neural activity. Other studies have used intracortical microarrays to decode speech^35–38^ or a neurotrophic electrode^39^ to synthesize formant frequencies^40, 41^ from the motor cortex. An alternative measure of intracranial neural activity is stereotactic EEG (sEEG), in which electrode shafts are implanted into the brain through small burr holes^42^. sEEG is considered to be minimally invasive, as a large craniotomy is not necessary and the infection risk is therefore smaller^43^. Additionally, the method of implanting the electrodes is very similar to that used in Deep Brain Stimulation (DBS), a method that has been used in the treatment of Parkinson’s Disease for several decades. In DBS, electrodes routinely remain implanted for many years, giving hope for the potential of sEEG for long-term BCIs. Similar to ECoG, sEEG is used in the monitoring of epilogenic zones in the treatment of refractory epilepsy. Between 5 and 15 electrode shafts are typically implanted covering a large variety of cortical and sub-cortical brain areas. Here lies one of the main differences to ECoG: instead of high density coverage of specific regions, sEEG provides sparse sampling of multiple regions. This sparse sampling could provide great potential for various BCI applications^44^.

Invasive recordings of neural activity are usually obtained during surgeries for seizure localization or glioma resection and are therefore not available to many researchers working on traditional speech decoding or speech synthesis technologies. These researchers are part of an active research community investigating the potential of non-invasive brain measurement technologies for speech neuroprostheses. Techniques include scalp-electroencephalography (EEG)^45–51^, providing high temporal resolution, especially the Kara One database^52^ provides the foundation for many studies; Magnetoencephalography^53, 54^, providing more localized information than EEG; and Functional Near Infrared Spectroscopy^55–58^, providing localized information of cortical hemoglobin levels. The advances made by this community could also benefit invasive speech neuroprostheses, a dataset that is provided to everyone could be used to evaluate and leverage their approaches.

To facilitate an increased understanding of the speech production process in the brain, including deeper brain structures, and to accelerate the development of speech neuroprostheses, we provide this dataset of 10 participants speaking prompted words aloud while audio and intracranial EEG data are recorded simultaneously (Fig. 1).

**Figure 1.**
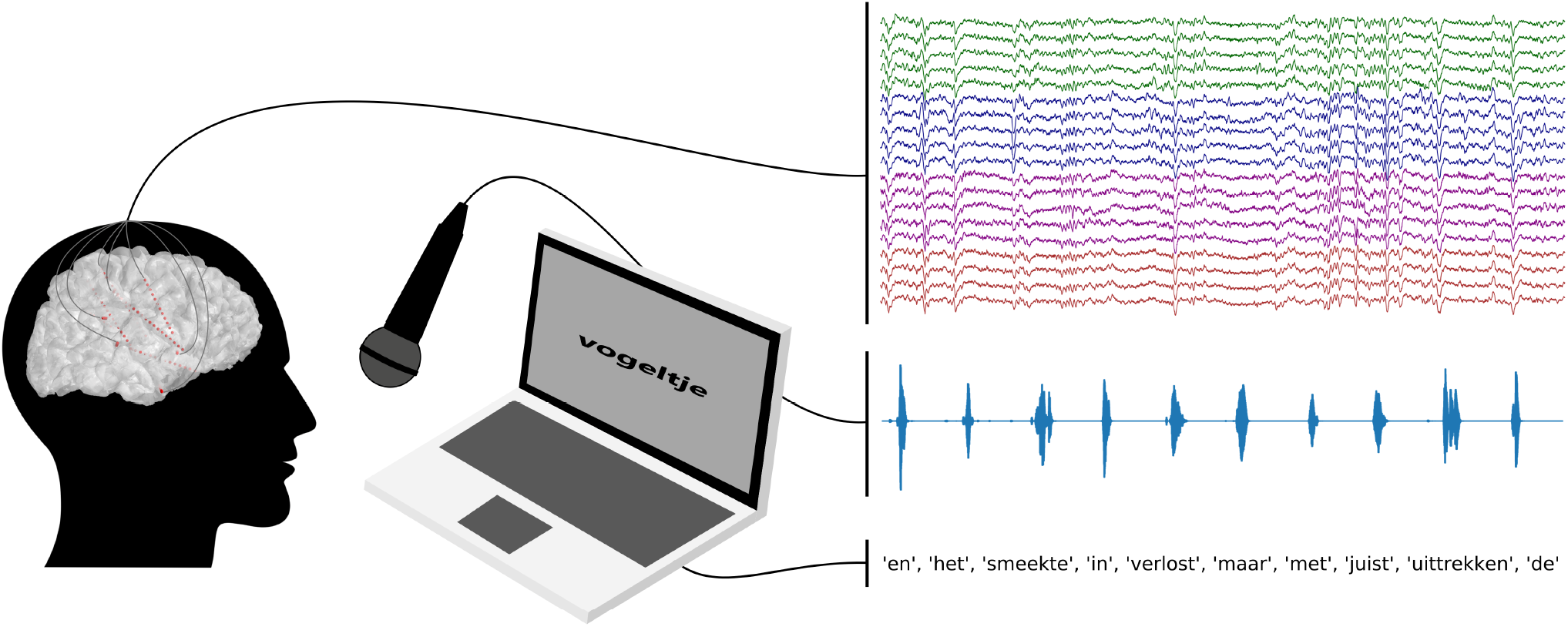
Intracranial EEG and acoustic data are recorded simultaneously while participants read Dutch words shown on a laptop screen. Traces on the right of the figure represent 30 seconds of iEEG, audio and stimulus data. The colors in the iEEG traces represent different electrode shafts.

## Methods

### Participants

A total of 10 participants suffering from pharmaco-resistant epilepsy participated in our experiment (mean age 32.4 ±12.6 years; 5 male, 5 female). Patients were implanted with sEEG electrodes as part of the clinical therapy for their epilepsy. Electrode locations were purely determined based on clinical necessity. All participants joined the study on a voluntary basis and gave written informed consent. Experiment design and data recording were approved by the IRBs of both Maastricht Unviversity and Epilepsy Center Kempenhaeghe. Data recording was conducted under the supervision of experienced healthcare staff. All participants were native speakers of Dutch. Participants’ voices were pitch-shifted to ensure anonymity.

### Experimental Design

In this study, participants were asked to read aloud words that were shown to them on a laptop screen (Fig. 1). A random word from the dutch IFA corpus^59^ and the numbers 1 to 10 was shown for 2 seconds during which patients read the word aloud. After each word, a fixation cross was displayed for one second. A total of 100 words was recorded for each participant resulting in a total recording time of 300 seconds.

### Data Acquisition

Participants were implanted with platinum-iridium sEEG electrode shafts (Microdeep intracerebral electrodes; Dixi Medical, Beçanson, France) with a diameter of 0.8 mm, a contact length of 2 mm and a inter-contact distance of 1.5 mm. Each electrode shaft contained between 8 and 18 electrode contacts. Neural data was recorded using one or two Micromed SD LTM amplifier(s) (Micromed S.p.A., Treviso, Italy) with 128 channels each. Electrode contacts were referenced to a common white matter contact. Data were recorded at either 1024 Hz or 2048 Hz and subsequently downsampled to 1024 Hz. We used the onboard microphone of the recording notebook (HP Probook) to record audio at 48 kHz. Audio data was subsequently pitch-shifted to ensure our participants’ anonymity using LibRosa^60^. We used LabStreamingLayer^61^ to synchronize audio and neural data and the experiment markers.

### Anatomical Labeling

Electrode locations (Fig. 2) were detected using the img_pipe Python package^62^ for anatomical labeling of intracranial electrodes. Within the package, for each participant, a pre-implantation anatomical T1-weighted Magnetic Resonance Imaging (MRI) scan was parcellated using Freesurfer (http://surfer.nmr.mgh.harvard.edu/), a post-implantation Computer Tomography (CT) scan was co-registered to the MRI scan and electrode contacts were manually localized. Their anatomical location label was extracted from the Destrieux atlas^63^ based parcellation.

**Figure 2.**
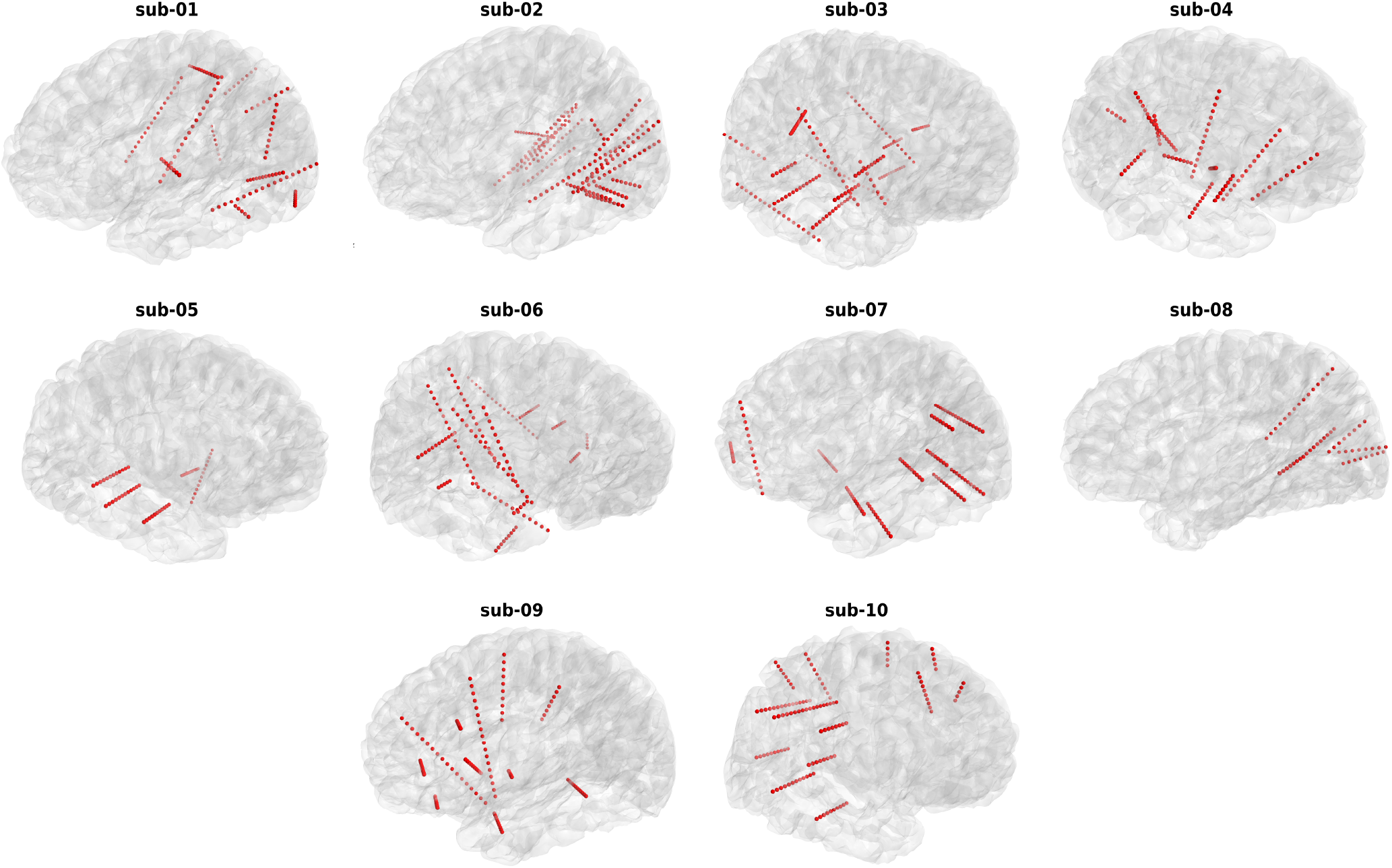
Electrode locations of each participant in the surface reconstruction of their native anatomical MRI. Each red sphere represents an implanted electrode channel.

By far the most electrodes are located in white matter (40.3%) and unknown areas (12.6%). Unknown areas are contacts that were not able to be labelled through the Freesurfer parcellation, for example a contact located just outside of the cortex. Thereafter, electrodes are predominantly located in the superior temporal sulcus, hippocampus and the inferior parietal gyrus.n See Fig. 3 for a full breakdown of anatomical regions and the number of electrodes implanted in those areas.

**Figure 3.**
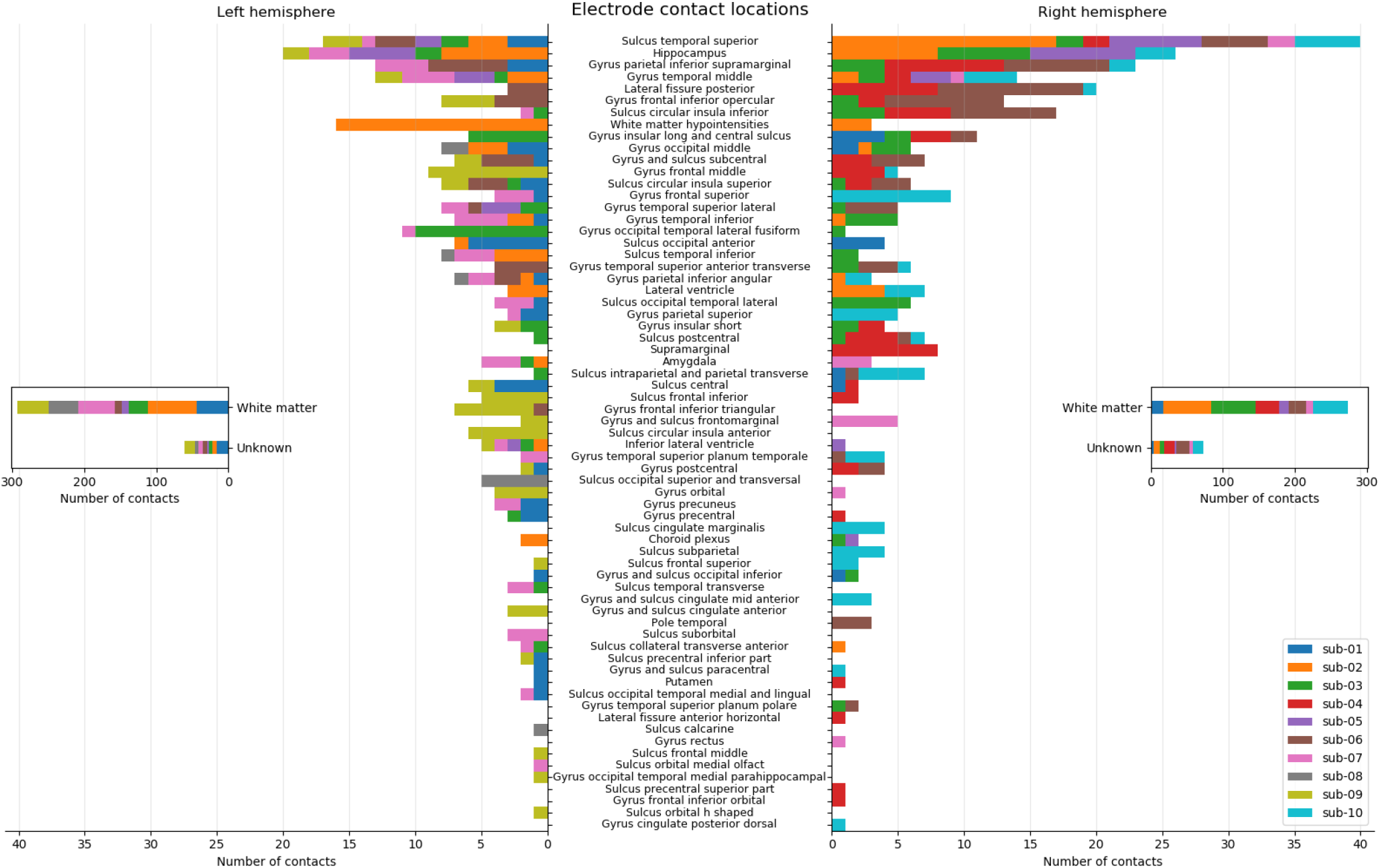
Number of electrode contacts in cortical and subcortical areas across all participants. Colors indicate participants. Lengths of the bars show the number of electrodes in the specified region. Note the deviant x-axis for the white matter and unknown regions.

### Data Records

The dataset is available at https://osf.io/nrgx6/. The raw data files (XDF format) were converted to Neurodata Without Borders (NWB; https://www.nwb.org/) format and organised in the iBIDS^64^ data structure format using custom Python scripts. The NWB format allows for compact storage of multiple data streams within a single file. It is compatible to the iBIDS structure, a community-driven effort to improve the transparency, reusibility and reproducibility of iEEG data.

The data is structured following the BIDS version 1.7.0 specification (https://bids-specification.readthedocs.io/en/stable/). The root folder contains metadata of the participants (*participants*.*tsv*), subject specific data folders (i.e., *sub-01*) and a derivatives folder. The subject specific folders contain .tsv files with information about the implanted electrode coordinates (*_electrodes*.*tsv*), recording montage (*_channels*.*tsv*) and event markers (*_events*.*tsv*). The *_ieeg*.*nwb* file contains three raw data streams as timeseries (iEEG, audio and stimulus). Descriptions of recording aspects and of specific .tsv columns are provided in correspondingly named .json files (i.e., *participants*.*json*). The derivatives folder contains the pial surface cortical meshes of the right (*_rh_pial*.*mat*) and left (*_lh_pial*.*mat*) hemisphere per subject derived from the Freesurfer pipeline. The meshes are provided for plotting utilities rather than the T1-weighted anatomical scan, to retain the anonymity of the participants. The description column in the *_channels*.*tsv* file refers to the anatomical labels derived from the Destrieux atlas^63^.

The iBIDS dataset passed a validation check using the BIDS Validator (https://bids-standard.github.io/bids-validator/) and manual inspection of each datafile.

### Technical Validation

We validate the recorded data by demonstrating that a spectral representation of speech can be reconstructed from the neural recordings using a simple linear regression model. This analysis is similar to a previous analysis in ECoG^65^.

### Checking for Acoustic Contamination

Acoustic contamination of neural recordings has been reported by Roussel et al.^66^. To ensure that the presented dataset does not contain acoustic contamination in the neural timeseries, we apply the method provided by the authors and correlate spectral energy between audio and neural data. We do not find any significant correlations (*p >* 0.01) on the diagonal of the contamination matrix for any of the participants.

### Feature Extraction

We extract the Hilbert envelope of the high-gamma band (70-170 Hz) for each contact using an IIR bandpass filter (filter order 4). To attenuate the first two harmonics of the 50 Hz line noise, we used two IIR bandstop filters (filter order 4). We averaged the envelope over 50 ms windows with a frameshift of 10 ms. To include temporal information into the decoding process, non-overlapping neighboring windows up to 200 ms into the past and future were stacked. Features are normalized to zero mean and unit variance using the mean and standard deviation of the training data. The same transform is then applied to the testing data.

The audio data is first downsampled to 16 kHz. To extract features for the audio data, we subsequently calculated the Short-Term-Fourier-Transform in windows of 50 ms with an frameshift of 10 ms. As the frameshift between neural and audio data is the same, there is a correspondence between audio and neural feature vectors. The resulting spectrogram is then compressed into a log-mel representation^67^ using 23 triangular filter banks.

### Decoding Model

To reduce the dimensionality of our decoding problem, we compress the feature space to the first 50 principal components. Principal components are estimated for each fold individually on the training data. We reconstruct the log-mel spectrogram from the high-gamma features using linear regression models. In these models, the high-gamma feature vector is multiplied with a weight matrix to reconstruct the log-mel spectrogram. The weights are determined using a least-squares approach.

### Waveform Reconstruction

The log-mel spectrogram does not contain the phase information anymore and an audio waveform can thus not be reconstructed directly. We utilize the method by Griffin and Lim^68^ for waveform reconstruction, in which the phase is initialized with noise and then iteratively modified. For a good algorithmic discription of the method, see^69^.

## Results

All results are obtained in a non-shuffled 10-fold cross validation in which 9 folds are used for training and the remaining fold is used for evaluation. This process is repeated until each fold has been used for evaluation exactly once.

### Spectrograms can be reconstructed

We evaluate the spectral reconstructions in terms of Pearson correlation coefficient between the spectral coefficients of the original speech and the reconstructed speech. For all 10 participants, speech spectrograms can be reconstructed from the neural data using linear regression (Fig. 4a). Reconstruction results were consistent across all 23 mel-scales spectral coefficients (Fig. 4b). Inspecting the spectrograms further (Fig. 5a), it can be seen that the results are mostly driven by the accurate reconstruction of speech versus silence. The spectral variations within speech are not captured by the linear regression approach. The Pearson correlation is not a perfect evaluation metric as this lack of detail during speech does not have a large impact on the score. We utilize the Pearson correlation here as a better metric has yet to be identified. By providing this open dataset,we hope that researchers developing more advanced metrics, such as the Spectro-Temporal Glimpsing Index (STGI)^70^ or the extended Short-Time Objective Intelligibility (eSTOI)^71^, will have the means to address this problem. For better reconstruction of speech, models that are more informed about speech processes (e.g. Unit Selection^28^) or neural network approaches with enough trainable parameters to produce high-quality speech^25, 26, 72–74^ might be necessary.

**Figure 4.**
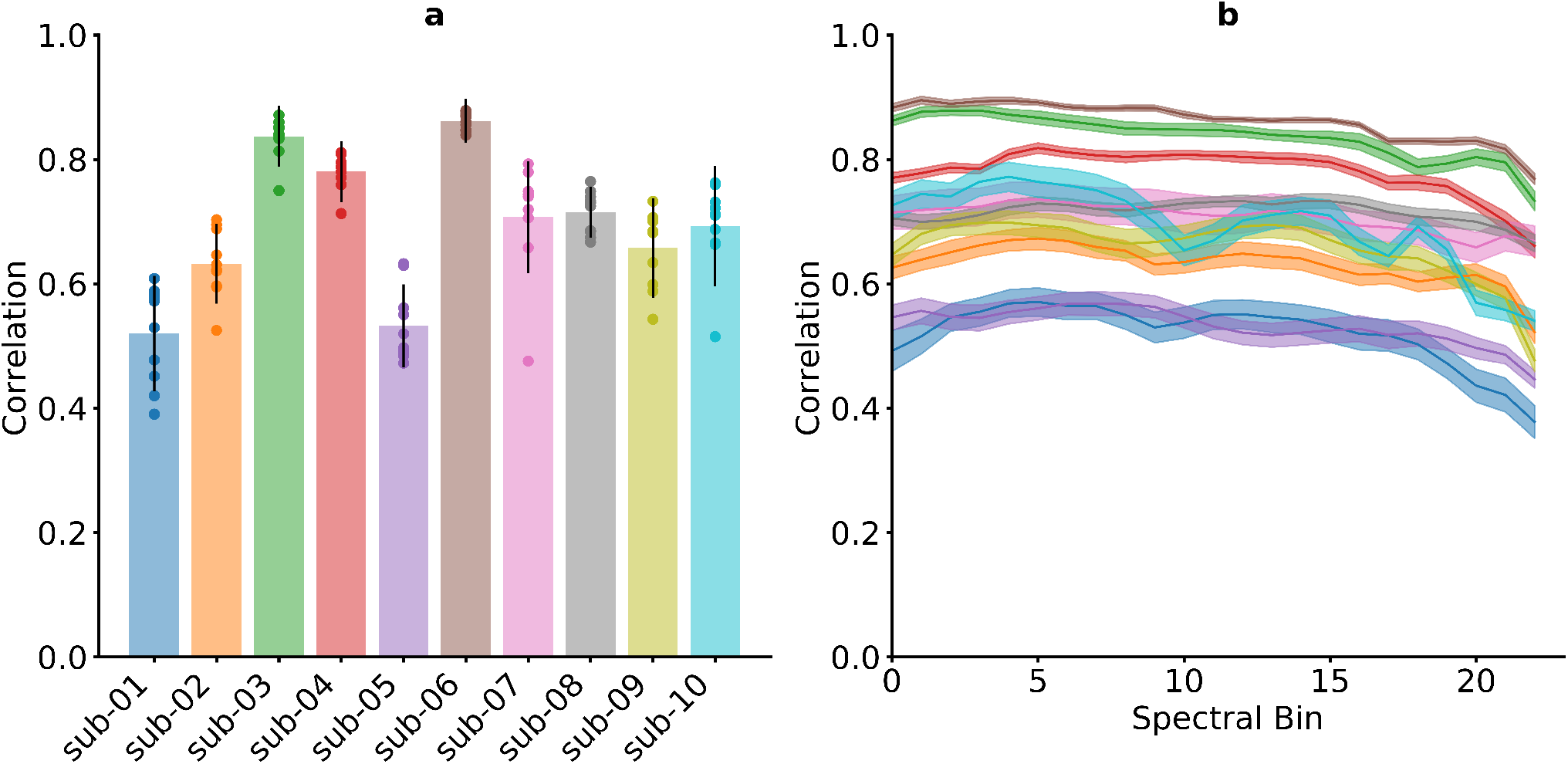
Results for the spectral reconstruction. (**a**) Mean correlation coefficients for each participant across all spectral bins and folds. Reconstruction of the spectrogram is possible for all 10 participants. Whiskers indicate standard deviations. Results of individual folds are illustrated by points. (**b**) Mean correlation coefficients for each spectral bin. Correlations are stable across all spectral bins. Shaded areas show standard errors.

**Figure 5.**
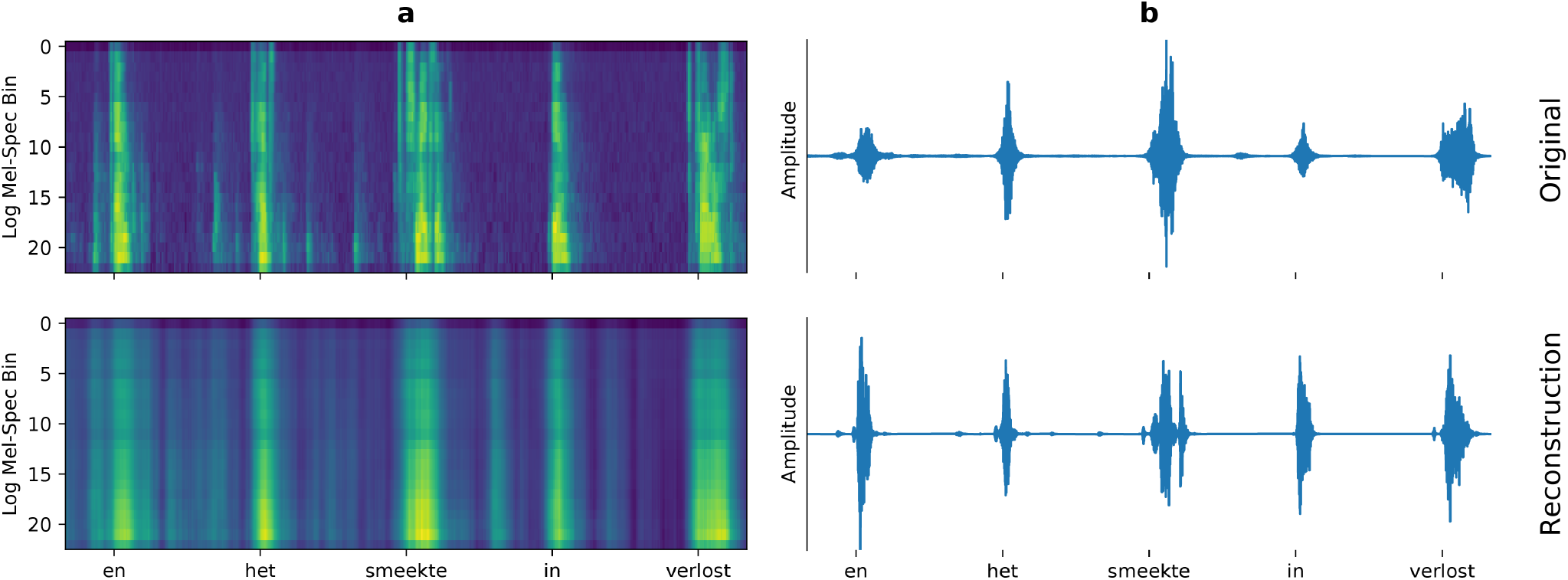
Spectrograms (**a**) and waveforms (**b**) of the original (top) and reconstructed (bottom) audio. The example contains five individual words from sub-06. While the linear regression approach captures speech and silent intervals very accurately, the finer spectral dynamics within speech are lost.

### Waveforms of Speech can be Reconstructed

Using the method by Griffin and Lim^68^, we can recreate an audio waveform from the reconstructed spectrograms (Fig. 5b). The timing between original and reconstructed waveforms is very similar, but listening to the audio shows that a substantial part of the audio quality is lost due to the synthesis approach. This is particularly clear when listening to the audio recreated from the original spectrogams (*_orig_synthesized*.*wav*). State-of-the-art synthesis approaches, such as WaveGlow^75^ or WaveNet^76^, could lead to better quality reconstructions.

## Usage Notes

### iBIDS data

Scripts to handle the data can be obtained from our github repository.

### Data loading and feature extraction

Neural traces, synchronized audio and experimental markers can be loaded using the provided *extract_features*.*py* script. High-gamma features are subsequently extracted and aligned to logarithmic mel-scaled spectrograms. Electrode channel names are also loaded.

### Spectrogram Reconstruction & Waveform Synthesis

Reconstruction of the spectrogram as well as the resynthesis to an audio waveform is performed in the *reconstruction_minimal*.*py* script.

### Anatomical data

Electrode locations can be found in the participant folder (*_electrodes*.*tsv*) and then be visualized using the cortical meshes (*_lh_pial*.*mat* and *_rh_pial*.*mat*) within the derivatives folder. These mesh files contain vertices coordinates and triangles, which are described by indices corresponding to vertex numbers.

## Code availability

All Python code to re-run the technical validation described in this report can be found on our github. The code relies on the numpy^77^, scipy^78^, pynwb^79^, scikit-learn^80^ and pandas^81^ packages.

## Acknowledgements

CH acknowledges funding by the Dutch Research Council (NWO) for the project ‘DEcoding Speech In SEEG (DESIS)’.

## Author contributions statement

C.H., P.L.K., A.J.C., L.W. and S.T. designed the experiment. M.V., M.C.O. and C.H. recorded the data. M.V. and S.G. localized the electrodes. M.V and C.H. analyzed the data and wrote the manuscript. All authors approve of the manuscript.

